# Coordinated organic and inorganic nitrogen transformations fuel soil microbial blooms and increase nitrogen retention during snowmelt

**DOI:** 10.1101/2024.11.30.626193

**Authors:** Patrick O. Sorensen, Ulas Karaoz, Harry R. Beller, Markus Bill, Nicholas J. Bouskill, Jillian F. Banfied, Rosalie K. Chu, David W. Hoyt, Elizabeth Eder, Emiley Eloe-Fadrosh, Allison Sharrar, Malak M. Tfaily, Jason Toyoda, Nikola Tolic, Shi Wang, Allison Wong, Kenneth H. Williams, Yangquanwei Zhong, Eoin L. Brodie

## Abstract

Snowmelt in high-elevation watersheds triggers a microbial bloom and crash that affects nitrogen (N) export. Predicting watershed N dynamics as snowpack declines is a challenge because the mechanisms that underlie this microbial bloom and crash are uncertain. Using a multi-omic approach, we show that the dynamic molecular properties of dissolved organic N, plus high gene expression for peptidases that recycle microbial biomass, suggested that microbial turnover provided N for biosynthesis during the microbial bloom. Amino acid fermentation by *Bradyrhizobia* produced organic acids that also fueled denitrification and dissimilatory nitrate reduction to ammonia (DNRA) during snowmelt. Nitrification in spring was driven by Spring-Adapted *Nitrososphaerales*, which utilized ammonium derived from osmolyte degradation by Winter-Adapted *Solirubrobacteraceae*. High DNRA gene expression after snowmelt suggested significant nitrate retention, increasing watershed N retention potential. However, declining snowfall may compromise microbial regulation of soil N retention, with implications for watershed N export.

## Introduction

Snowmelt is a biogeochemical hot moment that accounts for the largest annual river exports of nitrogen (N) in high-elevation watersheds^1,2^. Snowmelt also marks the transition from winter to spring and is associated with seasonal changes in solar radiation, warmer soil temperatures and higher plant primary productivity^3–6^. These environmental cues also initiate seasonal transitions for the soil microbiome^7–11^. For example, a substantial microbial bloom immobilizes soil N during snowmelt, which is followed by a biomass crash and pulse of soil N in ecosystems that have a seasonal snowpack^8,12–14^. Even though this phenomenon has been observed across a variety of ecosystem types^8,12–14^, the substrates and metabolic processes that fuel this soil microbial bloom are uncertain. The inorganic N that is exported from river discharge after snowmelt is primarily nitrate with an isotopic signature derived from soil microbial processes^15^. This implies that the snowmelt-associated microbial bloom and crash contribute to ecosystem N export. Higher air temperatures are reducing winter snowpack globally^16^, altering the seasonal timing, rate and magnitude of snowmelt^17^, and leading to altered watershed function in high-elevation catchments^18^. Currently, predicting how altered snowpack extent will affect watershed N export is challenging, in part because the microbial processes underlying the snowmelt-associated microbial bloom and crash are not well known.

Globally, soils turnover ∼240 Tg N year^-1^, with the majority of soil N being high molecular weight organic N^19,20^. Degradation of polymeric organic N is well-accepted as the rate-limiting step in soil N mobilization^21^, but despite this recognition, microbial N-transformations are often simplified to a few redox reactions involving mainly inorganic N species^22–24^, with organic N transformation either overlooked or only implicitly represented in models of the microbial N cycle. This is partly because the organic N pool is structurally diverse and has been difficult to assess until recently^25,26^. The uptake, assimilation, and mineralization of organic N are also coordinated by diverse metabolic systems that are regulated by different environmental signals^27–30^. Thus, these processes are emergent properties of microbiomes and challenging to ascribe to discrete taxa in complex communities. Whereas model organisms and *in vitro* studies have been used historically to describe microbial processes that underlie N degradation and assimilation^31,32^, recent metagenomic and metatranscriptomic analyses have also revealed the environmental diversity and distributions of organic N cycling processes in natural soils^33–35^. Simple classifications of microbial taxa into a small number of N cycling functional groups is likely insufficient because N-transforming taxa affect organic and inorganic N pools simultaneously^36^. Moreover, identifying interactions among organic and inorganic transforming taxa would lead to a more complete understanding of the microbial N cycle^37^.

Here, focusing on the snowmelt period within a headwater catchment of the Upper Colorado River Basin^2^, we combined genome-resolved metagenomics and metatranscriptomics to identify the metabolic pathways and processes that mobilize soil N during and after snowmelt. We also utilized high-resolution spectroscopy (Fourier Transform Ion Cyclotron Resonance Mass Spectrometry, FT-ICR MS; ^1^H nuclear magnetic resonance, NMR) to characterize low molecular weight N metabolites, the molecular properties, and transformations of the dissolved organic N (DON) pool. Snowmelt was previously shown to separate three distinct soil niches that are occupied by ecologically distinct microorganisms^12^. In this study, we hypothesized that these soil niches created by snowmelt are related to distinct metabolic capacities for degrading, assimilating and utilizing different N sources for growth and energy. Integrating multi-omic data sets, we found that ecologically distinct taxa are linked across the snowmelt time-period by their unique N cycling activities. These findings advance our understanding of microbial nitrogen cycling in high-elevation watersheds and have broader implications for predicting ecosystem responses to changing snowmelt patterns due to global change.

## Results

### The Soil Microbiome is the Critical N Control Point Before and After Snowmelt

Microbial biomass N was the largest extractable N pool and comprised 66% to 77% of the sum of all extractable soil N across all sampling dates (Figure 1a-c). Snow accumulation and snowmelt also affected the size and the relative proportions of most other soil N pools (Figure 1). For example, microbial biomass N and dissolved organic N (DON) increased 2.5-fold and 2-fold, respectively, during snowmelt in May as compared to maximum winter snowpack depth in March (Figure 1d,f). The microbial biomass pool size declined by 70% after soils became snow-free in June (Figure 1d) and DON followed a similar decline over that same time-period (Figure 1f). After the microbial biomass crash, we observed a pulse of inorganic soil N (Figure 1h, i), that included a 2-fold increase in ammonium and a 3-fold increase in nitrate. Soil sampling depth was not a significant factor influencing the response of soil N pools to snowmelt. We hypothesized that the pulse of ammonium and nitrate after snowmelt originated from immobilized organic N that was released after the microbial biomass crash. To evaluate this hypothesis, we next identified the microorganisms, N-substrates, and metabolic processes that sustain the snowmelt-associated microbial bloom and crash.

**Figure 1.**
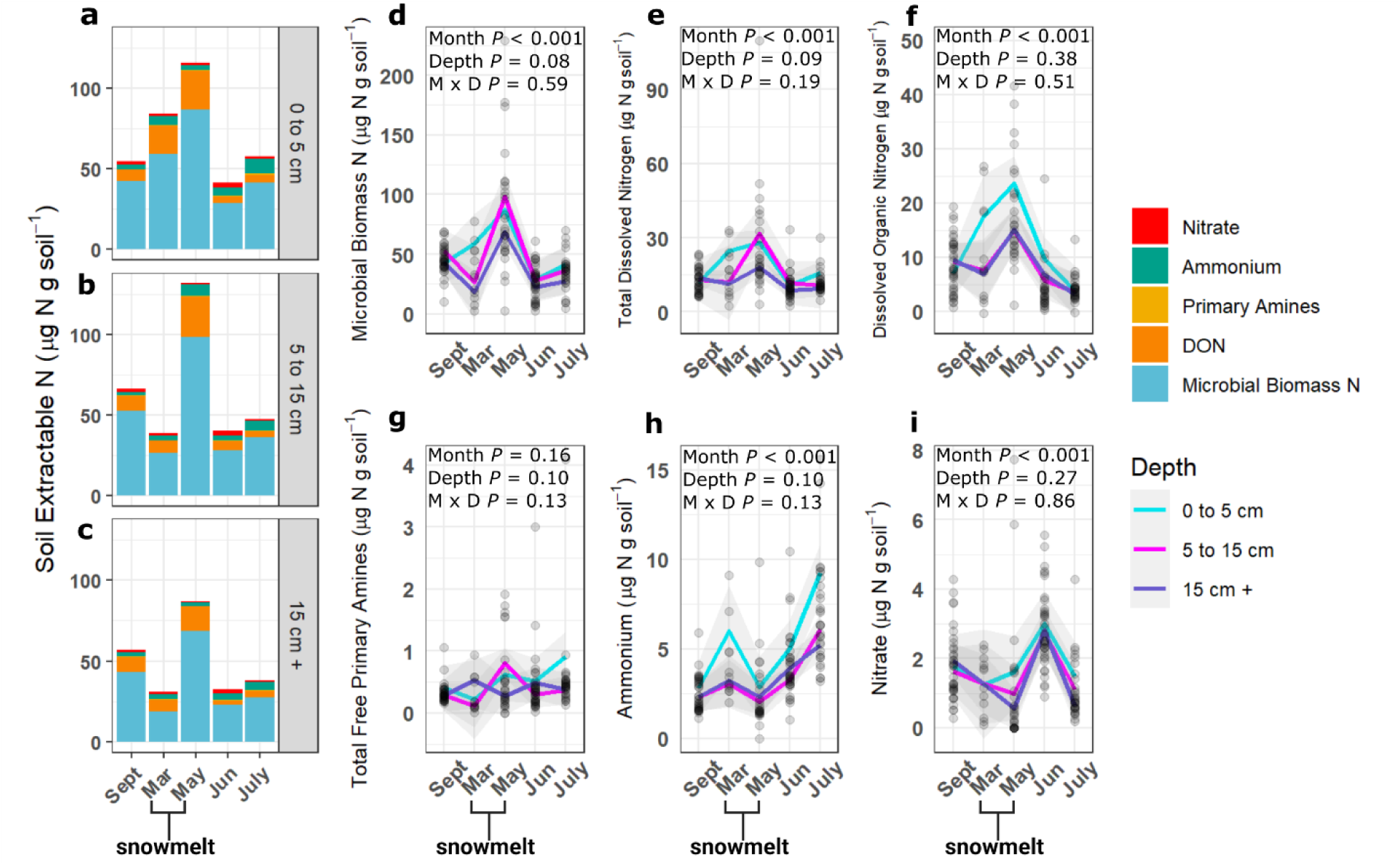
Soil extractable N pools across depths and time periods of winter snow accumulation (September to March), the snowmelt period (March to May) and loss of snowpack in spring (May to June) at East River, CO. The effects of month of sampling, depth and their interaction were tested using a mixed effects linear model (Figure 1d-i, n=12 on each date).

### Snow Accumulation and Snowmelt Control the Activity of Four Microbial Response Groups

We reconstructed 496 metagenome-assembled genomes (MAGs) across all sampling dates. A subset of 167 MAGs passed checkM with more than 75% completeness and less than 25% contamination (Supplemental Table 1). Winter snow accumulation followed by snowmelt imparted changes in microbial community composition and activity (Supplemental Figure 1). For example, the sampling month accounted for 18% of the variation in MAG genome coverage or 45% of the variation in MAG gene expression (both *P* <0.001, Supplemental Figure 1). Similar to the soil N pools, soil depth had a minimal effect on genome coverage or activity (Supplemental Figure 2).

Nearly all MAGs (139 out of 167 total high-quality MAGs) exhibited changes in total gene expression in response to snowmelt (Figure 2a). MAGs were clustered into microbial response groups based on MAG gene expression patterns across time (Figure 2a). The majority (77 out of 139 MAGs) were categorized as ‘Fall-Adapted’ and this response group collectively had more than a 2-fold increase in total gene expression after plant-senescence in September compared to peak snowpack depth in March. By contrast, ‘Winter-Adapted’ MAGs nearly doubled their total gene expression during the deepest winter snowpack in March compared to after plant senescence in September (Figure 2a). The most active Winter-Adapted MAGs constituted between 30% to 60% of total community gene expression in March and were dominated by a small number of organisms. This group included three *Bradyrhizobium* spp. MAGs (Alphaproteobacteria [Bin 167, Bin 164 and Bin 68]; 8%-20% total community expression), a single *Solirubrobacterales* MAG (Actinobacteriota [Bin 85]; 3% - 9% total community expression), and a single *Allosphingosinicella* MAG (Alphaproteobacteria [Bin 99]; 3% - 29% total community expression). Winter-Adapted MAGs had the highest GC content and the most active Winter-Adapted MAGs (e.g. Bins 167, 164, 99, 85) were amongst the lowest predicted optimal growth temperatures (Figure 2b), implying selection for growth underneath the snowpack during the coldest times of the year.

**Figure 2.**
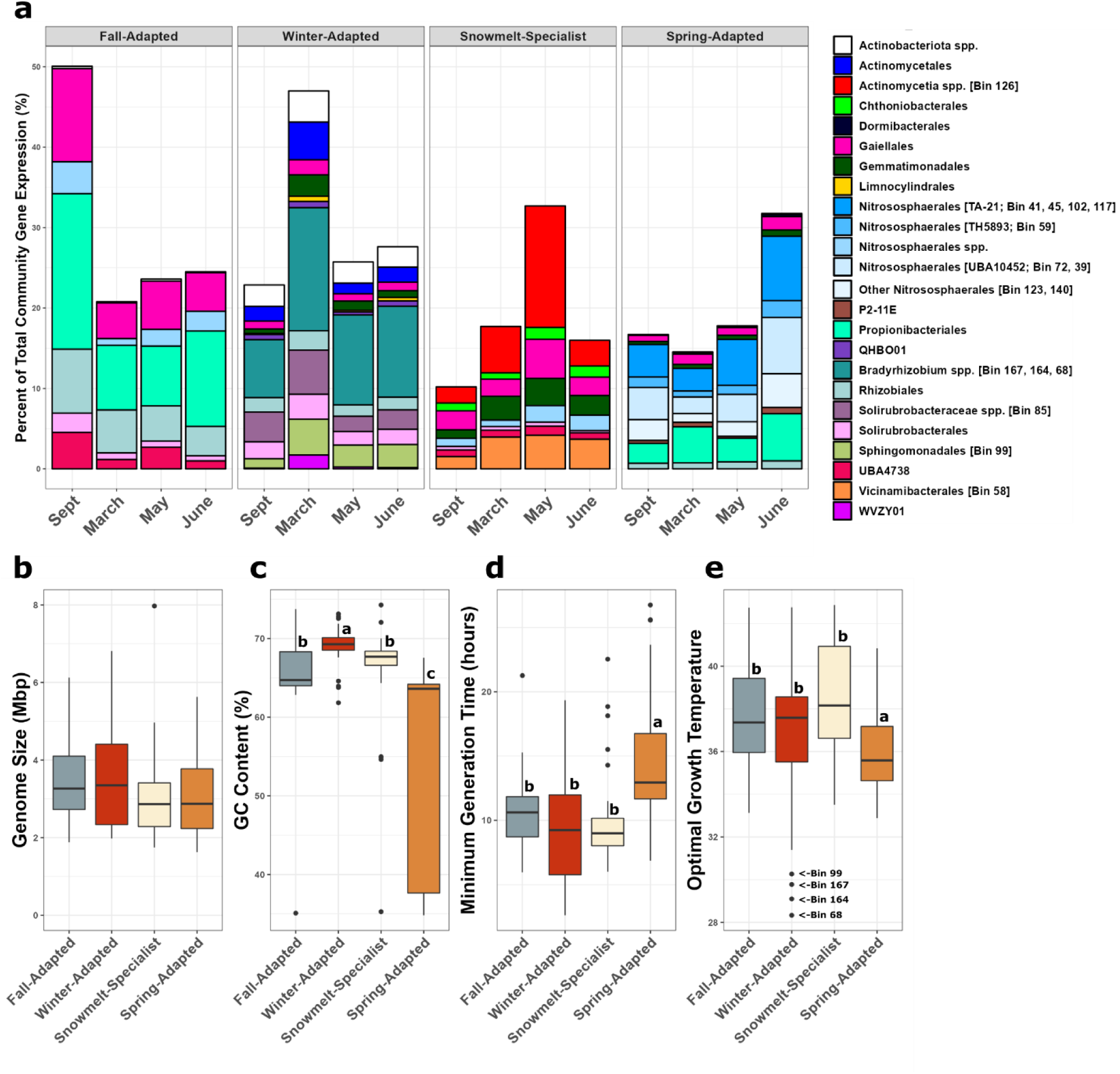
Metagenome-associated genomes (MAGs) were categorized into ecological response groups (Winter-Adapted, Snowmelt-Specialists, Spring-Adapted) using hierarchical clustering based on the date of each MAG’s maximum (a) RNA expression. MAG taxonomy is shown at the taxonomic Order level. MAGs with unresolvable taxonomy at the Order level were assigned a taxonomic affiliation based on the finest level of taxonomy that could be resolved. Differences in genomic traits across response groups (b-e) were tested using Dunn’s test and statistical differences are denoted with lower case letters (*P* ≤ 0.05).

Two ‘Snowmelt-Specialist’ MAGs (e.g. *Actinomycetia* spp. [Bin 126]; Acidobacteriota-Vicinamibacterales [Bin 58]) with high gene expression during snowmelt were disproportionately active relative to their abundance. For example, *Actinomycetia* spp. (Bin 126) accounted for 8% - 24% of total relative community gene expression in May, but only 1% - 2% relative abundance based on genome coverage (Supplemental Table 3). The ‘Spring-Adapted’ response group was dominated by 9 archaeal MAGs belonging to four genera (genus *UBA10452*, genus *TA-21*, genus *JAFAQB01*, genus TH56893; all Order Nitrososphaerales). These archaeal MAGs accounted for 23% of total community gene expression in June. Spring-Adapted organisms had the lowest GC content, longest predicted generation times, and low optimal growth temperatures (Figure 2c-e, Supplemental Table 1).

### Dissolved Organic N Became More Aromatic and Peptide-like During and After Snowmelt

Building on our analysis of microbial community dynamics, we next investigated how these microbial response groups degraded and were influenced by changes in dissolved organic nitrogen (DON). This approach allowed us to infer which N substrates fueled the microbial bloom during snowmelt. We observed 2700 to 4300 unique dissolved organic matter (DOM) compounds in each sample. Interestingly, the proportion of DOM compounds containing N (i.e. DON) varied significantly across the snowmelt period, comprising 8% of DOM compounds prior to the onset of winter versus 13% after the loss of snow cover in June (Supplemental Table 4). A shift in DON stoichiometry and molecular properties coincided with the changes in microbial community structure and activity (e.g. Figures 1,2). For example, the carbon to nitrogen ratio (C to N, Figure 3a), nominal oxidation state of carbon (NOSC, Figure 3b) and double bond equivalents (DBE, Figure 3d) within the DON pool were all highest prior to snowmelt compared to after snowpack loss. The reductions in NOSC and DBE that occurred during snowmelt implied the depletion of unsaturated, thermodynamically-favorable DON during the microbial bloom^38^, leading to the accumulation of more reduced and less thermodynamically favorable DON^39,40^. The aromaticity index of DON after plant senescence (Figure 3c) was indicative of highly-condensed, aromatic compounds^40^, then suggestive of a non-aromatic DON pool in March prior to snowmelt, followed by an increasingly aromatic DON pool during and after the microbial bloom in May (Figure 3c). Thus, the DON pool became more reduced, aromatic and accumulated N relative to C during and after snowmelt.

**Figure 3.**
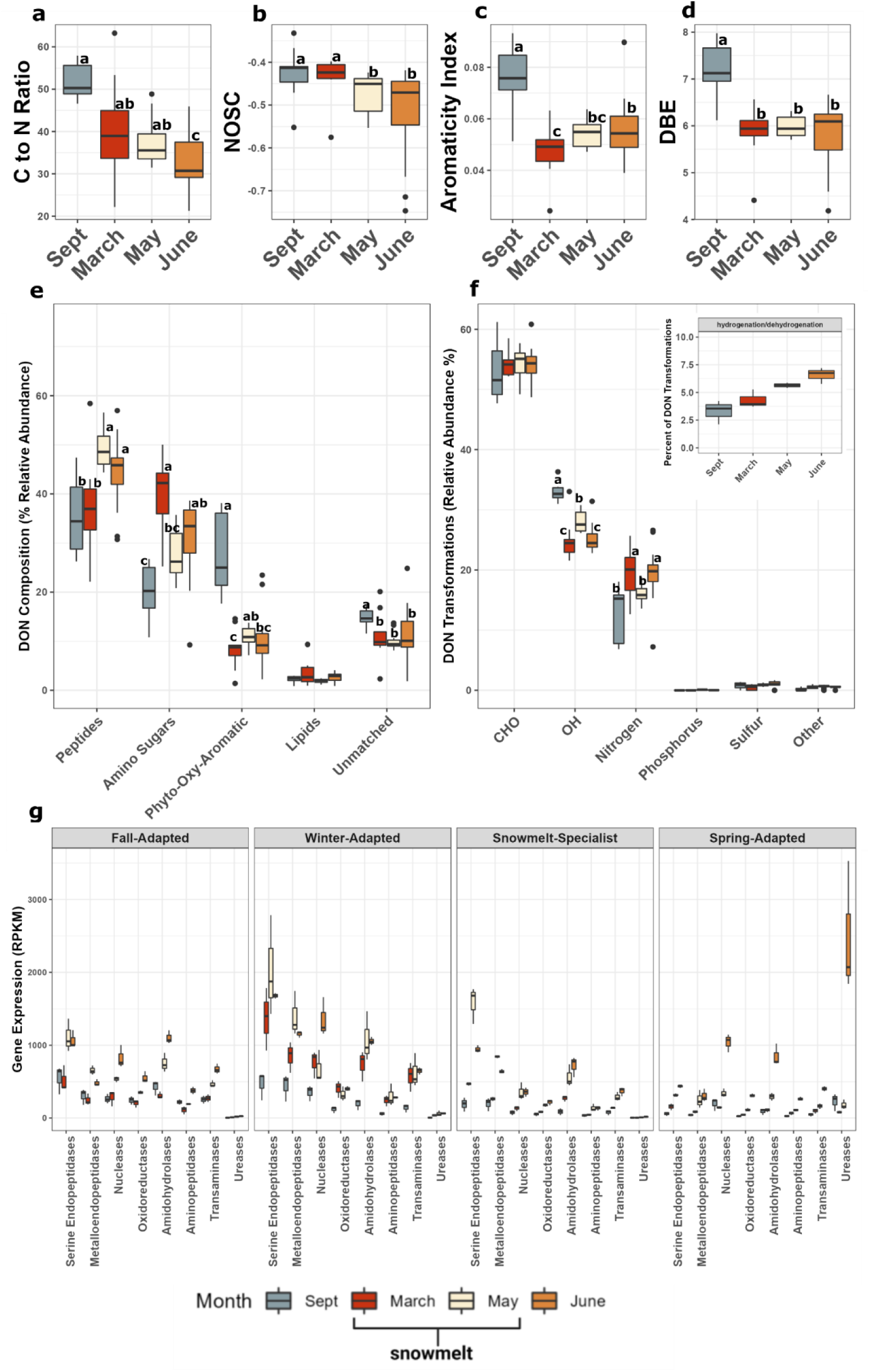
The molecular properties (a-b), composition of chemical classes (e) and transformation stoichiometric categories (f) of dissolved organic N (DON) detected by FT-ICR MS. The inset of (f) is the frequency of ‘hydrogenation/dehydrogenation’ transformations which were the most frequently detected of all transformations and a subset of ‘O/H’ transformations. The effect of month was tested using non-parametric Dunn’s tests. Differences between dates were determined post hoc and statistical differences are denoted by different letters (*P* ≤ 0.05). Gene expression of proteolytic enzymes (g) were summed by microbial response groups and difference in gene expression were tested non-parametric Dunn’s test with significant time effects denoted *** *P* ≤ 0.001, \*\**P* ≤ 0.01, \**P* ≤ 0.05

Peptide-like compounds were the most frequently detected chemical class of DON and accounted for 22% to 58% of all DON compounds (Figure 3e). The molecular composition and transformations of DON (Figure 3f) were also reflective of changes in DON stoichiometric and molecular properties (e.g. 3a-d). In this context, we define ‘transformations’ as chemical modifications resulting in the loss or gain of functional groups (e.g. O/H, or N-containing functional groups), which can significantly alter the molecular structure and bioavailability of DON compounds. Phyto-oxy-aromatic-like DON compounds were most frequently detected after plant senescence and declined two-fold between September and March (Figure 3e). The relative abundance of amino sugar-like compounds doubled during that same time (Figure 3e). We also observed up to 16-fold increases in gene expression for enzymes that degrade chitin (e.g. *chbG*) or recycle peptidoglycan (e.g. *nagA, B, Z; mepA,H; mrcA,B*) by Winter-Adapted MAGs from September to March. These patterns suggest that the initial source of DON pool was plant litter under snow, but that the degradation of microbial biomass became an important source of amino sugar-like compounds during winter and prior to snowmelt (Figure 3e).

The proportion of peptide-like compounds was 25% higher during snowmelt in May as compared to March and phyto-oxy-aromatic compounds followed a similar trend (Figure 3e). In addition, we detected 103 different transformations that resulted in the loss or addition of N-containing monomers (Supplemental Table 5). The frequency of these DON transformations declined (86 out of 103; including amino acids see Supplemental Figure 3) when microbial N immobilization was highest (Figure 1a,d). Concurrently, we observed between 4-fold to 30-fold higher gene expression for serine endopeptidases among Winter-Adapted and Snowmelt-Specialist MAGs during snowmelt as compared to before snowmelt ensued (Figure 3g). Metalloendopeptidases, amidohydrolases, and nucleases followed a similar trend. Transformations of DON involving the loss or addition of oxygen, hydrogen or their combination (e.g. “hydrogenation”, see Supplemental Table 5 “O/H”) increased significantly during snowmelt in May as well (Figure 3e). Collectively, these findings suggested that the DON pool was predominantly transformed by endopeptidase, amidohydrolase, and nuclease activity during snowmelt. These enzymatic activities drove ‘O/H’-type transformations, resulting in an increased proportion of peptide- and aromatic-like DON (Figure 3c,e,f). This shift in DON composition aligned with the changing metabolic activities, particularly among the Winter-Adapted and Snowmelt-Specialist microbial response groups (Figure 3g).

### Maximum Ammonia Assimilation is Offset from the Snowmelt Biomass Bloom

Having observed dynamic changes in the molecular properties of DON during and after snowmelt, we next investigated how these changes in DON affected microbial N assimilation strategies. Gene expression for N assimilation from ammonia through the glutamine synthetase – glutamate synthase (GS-GOGAT) pathway was strongly affected by snowmelt (Figure 3a), as were transcription regulators controlling organic N uptake. Nitrogen in most nucleotides is derived from the amide bond in glutamine while the primary amino group for biosynthesis of all amino acids originates from glutamate^41^. Thus, the production of glutamine and glutamate are central to cellular growth, and we expected the GS-GOGAT pathway to be most highly expressed during the microbial bloom (e.g. Figure 1). Glutamate showed the highest concentration of all small-molecule metabolites containing N detected across all dates (e.g. ∼20 nM to 3.4 mM; Figure 4b and Supplemental Table 6) and increased significantly between March and May (Figure 3a). As predicted, the temporal trend in glutamate production coincided with the temporal trend in microbial biomass production during snowmelt (see Figure 1a,e).

**Figure 4.**
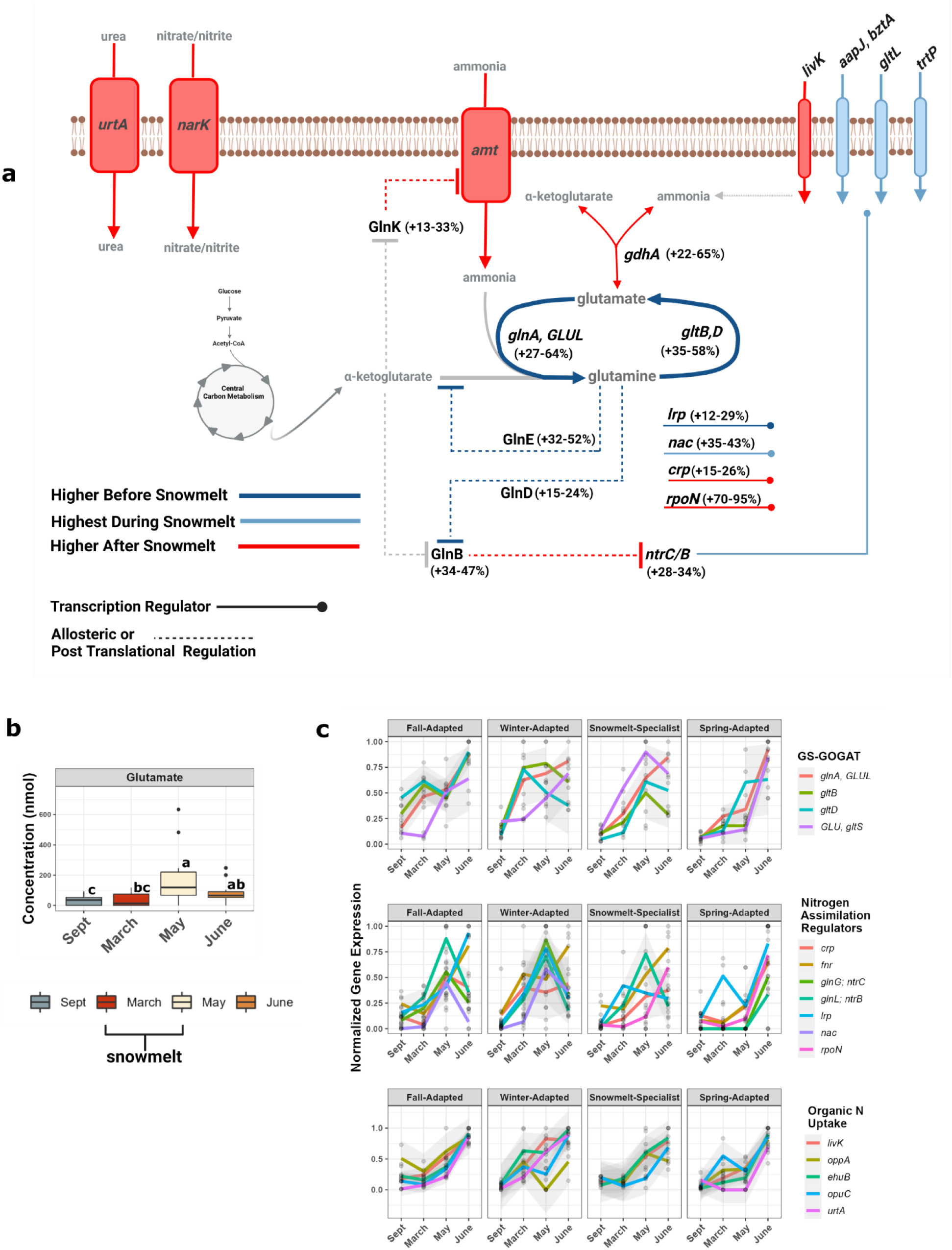
Gene expression at community-level (a) of key enzymes and transcription regulators of microbial N assimilation and glutamate production (b) across dates. Gene expression in (a) was categorized as ‘Higher Before Snowmelt’ (March relative to May gene expression; dark blue lines), ‘Highest During Snowmelt’ (May relative to June or March; light blue lines) or ‘Higher After Snowmelt’ (June relative May gene expression; red lines) with range derived from replicates sampled across soil depths. Gene products in (a) with allosteric binding or post-translation regulation capacity are shown upper-case and metabolites are depicted in gray text. In panel (c) gene expression was normalized (0-1) by the maximum gene expression for each gene within each microbial response group across dates. Abbreviations - glutamine synthase (*glnA, GLUL*), glutamate synthetase small and large subunit (*gltB,D*), glutamate dehydrogenase (*gdhA*), N regulatory PII-1 or PII-2 protein (GlnB or GlnK), glutamine synthetase adenylyltransferase (GlnE), glutamine synthetase uridylyltransferase (GlnD), NRII-NRI Two-Component Regulatory System (*ntrC/B*), cyclic AMP receptor protein (*crp*), leucine responsive protein (*lrp*), N assimilation control (*nac*), and RNA polymerase sigma factor N (*rpoN*; also known as σ^54^), branched-chain amino acid transport system substrate binding protein (*livK*), oligopeptide transport system substrate binding protein (*oppA*), ectoine transport system substrate binding protein (*ehuB*), choline/glycine betaine transport system substrate binding protein (*opuC*), urea transport system substrate binding protein (*urtA*).

However, total gene expression for the primary glutamine and glutamate producing enzymes (*glnA, GLUL*; *gltB,D*) were 27% to 58% higher under the winter snowpack in March compared to during snowmelt in May (Figure 4a), suggesting that ammonia assimilation via GS-GOGAT was offset from the biomass bloom that occurred during snowmelt (Figure 1a,d). Gene expression for the transcription factor leucine responsive protein (*lrp*; Figure 4a), and the post-translational, allosteric regulators of glutamine synthetase (GlnE – GlnD), followed the same trend. Lrp and glnE/D increase N assimilation from ammonia when glutamate and α-ketoglutarate intracellular concentrations are high^41^, both of which are indicated of high aerobic metabolism and high ammonium availability^42^ beneath the winter snowpack.

Maximum gene expression for N assimilation through the GS-GOGAT pathway was also different among microbial response groups (Figure 4c), implying niche partitioning of N assimilation related to snow accumulation and snowmelt. For example, Snowmelt-Specialist MAGs had maximum gene expression of GS-GOGAT genes during the microbial biomass bloom (Figure 4c), whereas maximum gene expression of GS-GOGAT genes occurred after snowpack loss for Spring-Adapted MAGs. Thus, maximum N assimilation through the GS-GOGAT pathway differed among response groups, corresponding with the period of maximum metabolic activity for each group.

In contrast to GS-GOGAT, transcription for the NRII-NRI Two-Component Regulatory System and N assimilation control transcription factor (*nac*) were overall more highly expressed during the snowmelt period in May (Figure 4a). Similar to GS-GOGAT genes, the timing for maximum gene expression of *ntrC/B* plus *nac* also differed among microbial response groups (Figure 4c). The NRII-NRI Two-Component Regulatory System is activated during N-limited growth marked by low external ammonia availability and low intracellular concentrations of glutamine^43^. Together, the NRII-NRI Two-Component System plus Nac form a regulon that controls the expression of more than 75 genes involved in utilizing organic N sources during periods of N-limited growth^44^. Winter-Adapted and Snowmelt-Specialist MAGs had maximum gene expression of the NRII-NRI Two-Component System plus Nac during the snowmelt period in May (Figure 4c), which coincided with increased expression for the uptake of branch-chained amino acids (*livK*) and ectoine/hydroxyectoine (*ehuB*). Altogether, these gene expression patterns support the hypothesis that uptake and degradation of organic N (e.g. peptide-like DON; see Figure 2a) fueled the microbial biomass bloom during snowmelt.

Gene expression after snowmelt suggested that the soil microbiome broadly switched from N- to C-limited growth at the start of the plant-growing season. For example, gene expression for ammonia assimilation via glutamine dehydrogenase (*gdhA*) increased by 22% to 65% after the loss of snow in June (Figure 4a), coinciding with a pulse of ammonium and nitrate (e.g. Figure 1d,h,i). Significant increases in expression of RNA polymerase σ^54^ (*rpoN*), the N regulatory PII-1 and PII-2 proteins (GlnB and GlnK), and the transcription factor cAMP receptor protein (*crp*) were also observed in June. The PII-1 and PII-2 proteins have RNA polymerase σ^54^ dependent promoters ^45,46^ and increase ammonia assimilation by inactivating NRII-NRI Two-Component System – Nac^47^. High gene expression of Crp is an indicator of low carbon to nitrogen substrate stoichiometry (see Figure 3), which activates carbon catabolite repression and increases the degradation of aromatic substrates^48–50^, consistent with increased potential for the degradation of aromatic DON observed after snowmelt (Figure 3e).

### Amino Acid Fermentation Fuels Bloom and Controls Fate of Nitrate During Snowmelt

We next quantified changes in low molecular weight N metabolites and compared these with gene expression for N cycling pathways among highly active MAGs to infer the fate of N during and after the microbial bloom and crash. High concentrations of branched-chain amino acids, including alanine, leucine, isoleucine and valine, were observed with α-keto acids that are one carbon atom shorter (e.g. pyruvate, 2-oxoisocaproic acid, isovalerate) during the microbial bloom. This co-occurrence suggests that amino acid fermentation (e.g. Strickland Oxidation reactions^51^) was an important pathway for N assimilation during snowmelt (Figure 5a). Transamination of amino acids, the first step in amino acid fermentation^52^, produces glutamate that can be used for biosynthesis, and we observed elevated transaminase gene expression during snowmelt in May (e.g. see Figure 4a).

**Figure 5.**
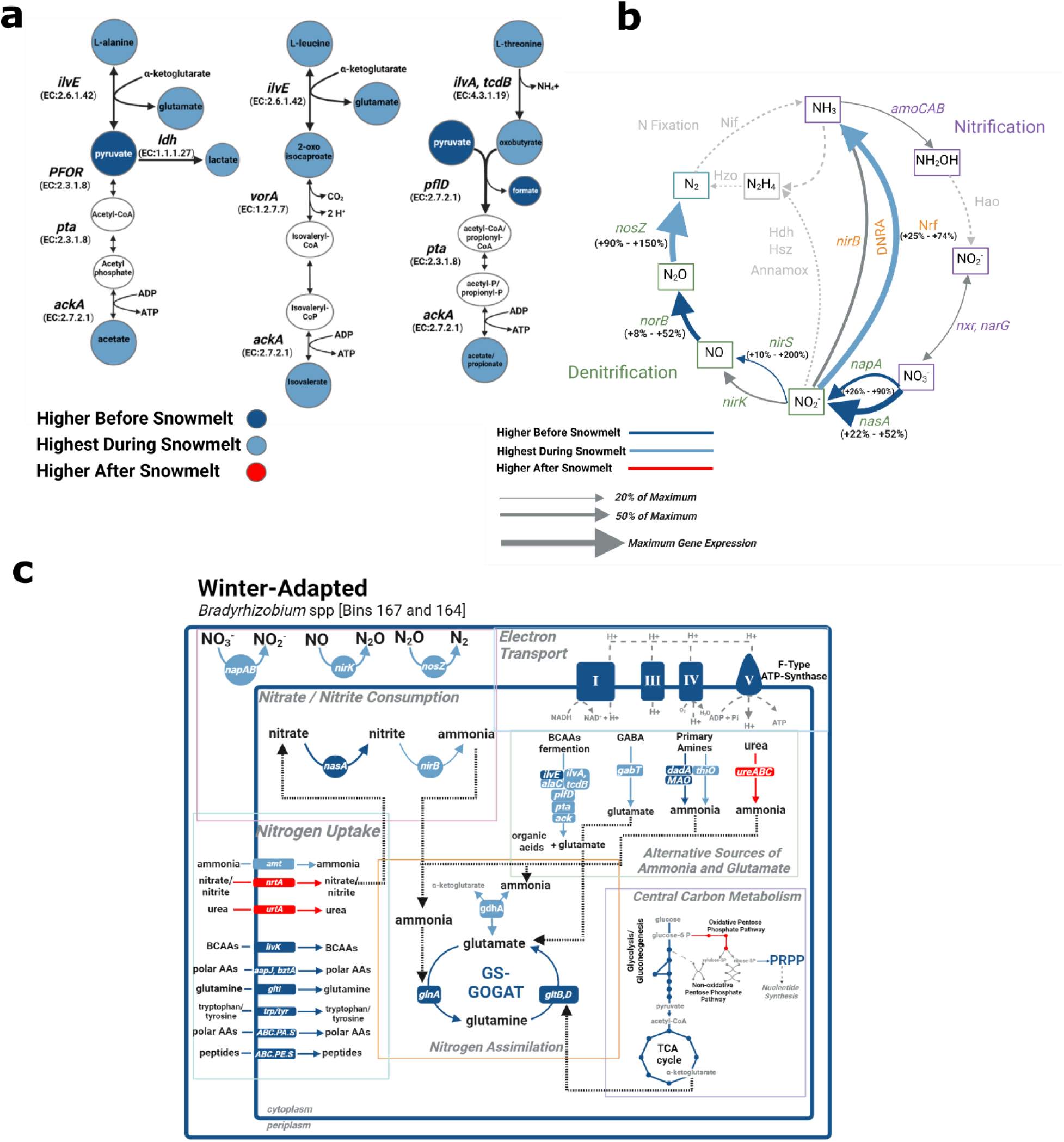
Fermentation of branched-chain amino acids (Strickland oxidative branch) that produces glutamate and organic acids was inferred from (a) small molecule metabolite concentrations detected by ^1^H-NMR. The color of filled compounds in (a) indicates the sampling date with highest compound concentration determined using Dunn’s test (*P* ≤ 0.05). Unfilled compounds were not detected in our dataset. Gene expression for (b) nitrate/nitrite consumption through denitrification or respiratory dissimilatory nitrate reduction to ammonia (DNRA) had their highest expression prior to or during snowmelt. Metabolic reconstruction of highly active Winter-Adaptive *Bradyrhizobium* spp. MAGs [Bins 167, 164] indicated (c) branched-chain amino acid fermentation coupled to during snowmelt to N_2_O consumption. Abbreviations for (a) expressed genes in the pathway are *ilvE* - branched-chain aminotransferase, *ilvA, tcdB* – threonine dehydratase, *ldh* – lactate dehydrogenase, *pflD* – pyruvate-formate lysase, *PFOR* – pyruvic-ferredoxin oxidoreductase, *vorA* - 2-oxoisovalerate ferredoxin oxidoreductase alpha subunit, *pta* - phosphate acetyltransferase, *ackA* – acetate kinase.

High accumulation of organic acids produced by amino acid fermentation likely affected nitrate consumption through competing pathways (e.g. denitrification versus dissimilatory nitrate reduction) that influence soil N retention versus loss (Figure 5b). For example, acetate, propionate and other products of amino acid fermentation (e.g. butyrate, H_2_) are electron donors for energy production^53^, and both denitrification and dissimilatory nitrate reduction to ammonia (DNRA) compete for nitrate as a terminal electron acceptor. DNRA is favored when non-fermentable, organic electron donors (e.g. acetate) are abundant and the ratio of reduced organic carbon to nitrate is high^54^, conditions that were generally observed during snowmelt (e.g. Figure 1e, 5a), and the highest gene expression for nitrate consumption through the respiratory branch of the DNRA pathway (e.g. *napAB* plus *nrfA*) occurred at that time (Figure 5b). By contrast, the highest potential for nitrite reduction to NO or N_2_O occurred underneath the deepest snowpack in March and prior to the accumulation of organic acids (e.g. Figure 5a). For example, nitrous oxide reductase (*nosZ*) gene expression was 90% to 150% higher during snowmelt in May as compared to March (Figure 4c). The denitrification-related gene expression patterns observed here are consistent with gas fluxes showing that soil N_2_O efflux was highest prior to the onset of snowmelt^55^, whereas the snowmelt period was a time of high N_2_O consumption.

Winter-Adapted *Bradyrhizobium* MAGs (e.g. Bins 164 and 167) were among the most active taxa underneath the winter snowpack in March (e.g. Figure 2a) and illustrated connections between amino acid fermentation and denitrification potential. For example, Winter-Adapted *Bradyrhizobium* spp. had high gene expression for amino acid and peptide uptake followed by deamination of amino acids underneath the winter snowpack (Figure 6c). Nitrogen assimilation through GS-GOGAT and ATP production via oxidative phosphorylation were also highest at that time. High gene expression for anaerobic degradation of amino acids in May (Figure 6c), suggested that Winter-Adapted *Bradyrhizobium* utilized glutamate produced via amino acid fermentation for N assimilation during snowmelt. Organic acids produced from amino acid fermentation may have been then used as electron donors for denitrification (Figure 5c), as evidenced by high gene expression for NO production and complete N_2_O reduction to N_2_. Collectively, the gene expression patterns observed indicate that these organisms utilize aerobic metabolism to produce energy and assimilate N as ammonia prior to snowmelt. During snowmelt, the Winter-Adapted *Bradyrhizobium* spp. fermented amino acids for N assimilation coupled to denitrification, highlighting this organism’s capacity for coordinated organic and inorganic N transformations for energy and growth during the microbial bloom.

**Figure 6.**
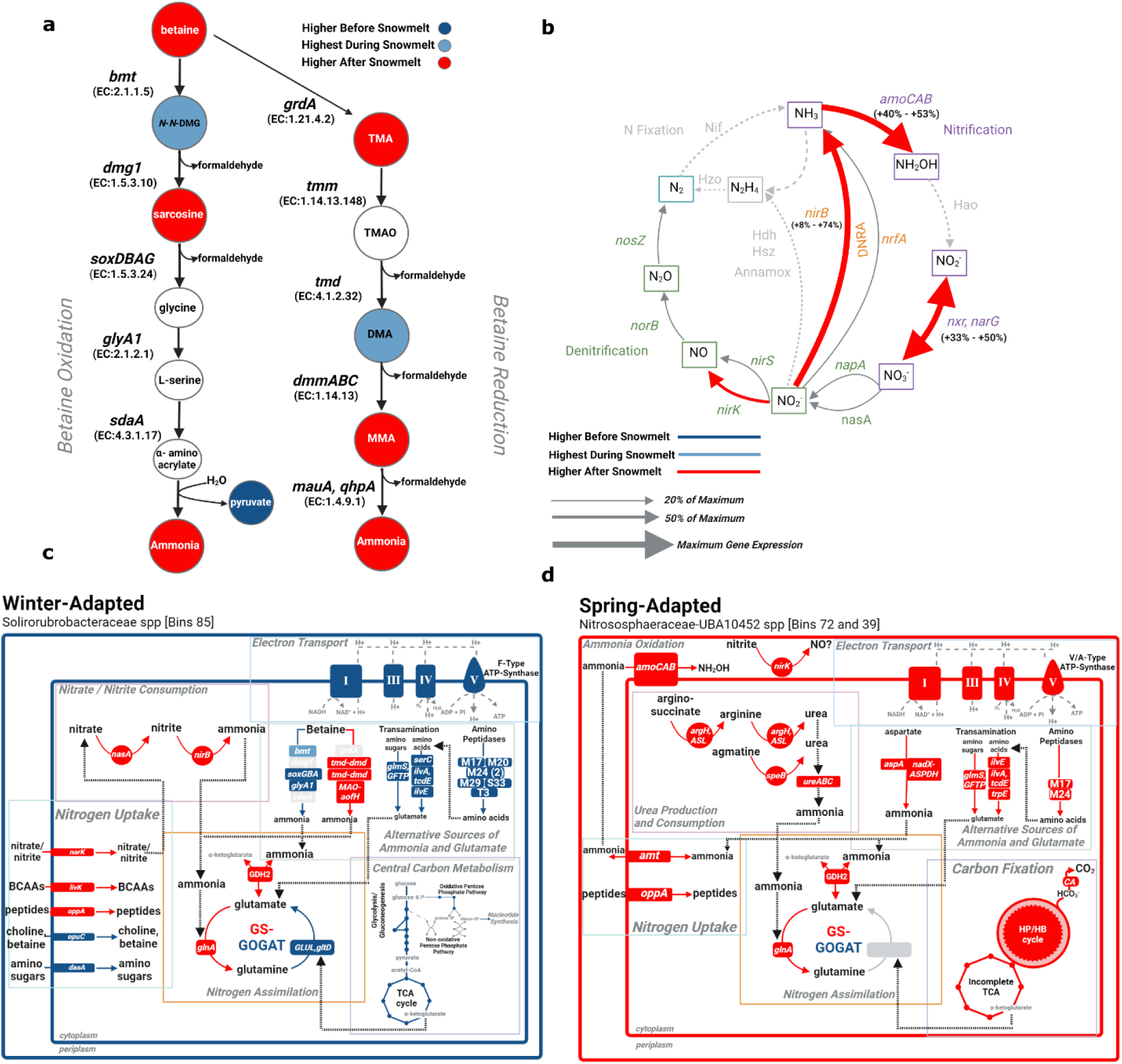
High concentrations of betaine and a suite of methylated amines indicated (a) active betaine oxidation or reduction before and after snowmelt, respectively. The color of filled compounds in (a) indicates the sampling date with highest compound concentration determined using Dunn’s test (*P* ≤ 0.05). Unfilled compounds were not detected in our dataset. Gene expression for (b) ammonia and nitrite oxidation (i.e. nitrification) were highest after the loss of snowpack in June as was fermentative DNRA. Metabolic reconstructions suggested that Winter-Adaptive Solirubrobacteraceae [Bin 85] actively degrades betaine after snowmelt which may provide N to Spring-Adapted Nitrososphaerales that actively oxidize ammonia and have high ureolytic activity. Abbreviations for genes in (a) are *bmt* – betaine-homocysteine S-methyltransferase, *dmg1*-dimethylglycine oxidase*, soxDBAG* – sarcosine oxidase, *glyA1* – serine dehydratase, *grdA* – betaine reductase, *tmd* – trimethylamine N-oxidase demethylase, *dmmABC* – dimethylamine monooxygenase, *mauA, qhpA* – methylamine dehydrogenase.

### Osmolytes Released from Biomass are a Source of Ammonia after Snowmelt

Betaine and an assortment of methylated amines had high concentrations after the biomass crash in June (Figure 6a). The observed methylated amines included degradation products characteristic of two different pathways of betaine catabolism: the oxidative demethylation pathway that is initiated by betaine-homocysteine methyltransferase^56^, or the degradation pathway that proceeds via trimethylamine and originates from a Strickland-like reduction of glycine betaine^57^. Genes related to the oxidative pathway of betaine degradation were more actively expressed prior to the onset of snowmelt in March whereas genes related to reductive betaine degradation were more actively expressed after snowmelt in June (Supplemental Figure 5). A highly active Winter-Adapted Solirubrobacteraceae MAG had the highest betaine degradation potentials of all organisms (Figure 6c). This MAG exhibited the highest gene expression for oxidative betaine degradation in March and high gene expression for reductive betaine degradation in June. Furthermore, this Winter-Adapted Solirubrobacteraceae MAG also demonstrated high aminopeptidase activity underneath the snowpack in March, producing amino acid monomers that were transaminated to produce glutamate at that time (Figure 6c). Thus, it is likely that this Winter-Adapted Solirubrobacteraceae MAG utilized ammonia from oxidative betaine degradation and glutamate from amino acid transamination as N sources for biosynthesis and growth underneath the winter snowpack.

The pulse of nitrate that we observed after snowmelt in June (e.g. Figure 1i) coincided with high gene expression of ammonia- and nitrite oxidation related genes. For example, genes associated with ammonia- or nitrite oxidation were 40% to 53% (*amoCAB;* Figure 6b) or 33% to 50% (*nxrA, narGH*; Figure 6b) more highly expressed after loss of snowpack in June as compared to May (Figure 6b). These patterns co-occurred with marked increases in the activity of ammonia-oxidizing Spring-Adapted Nitrososphaerales (Figure 6d; also Figure 2a). Spring-Adapted Nitrososphaerales had high capacity to produce urea from arginine or agmatine as well as urease activity that was up to 30-fold higher after snowpack loss. High gene expression for the cytoplasmic nitrite reductase (NADH-dependent) large and small subunits (*nirBD;* Figure 5b) were also 8% to 52% higher in June compared to during the snowmelt period in May. This indicated high potential for nitrate retention through the fermentative DNRA pathway after snowmelt, potentially limiting N leaching losses at that time.

## Discussion

Here, we integrated genome-resolved metagenomics, metatranscriptomics, and metabolomics to identify the sources of N that fuel the snowmelt-associated microbial bloom and to infer the fate of N after the microbial population crashes post-snowmelt. The stoichiometric and molecular properties of the dissolved organic N pool suggested that plant detritus was the initial source of N used for microbial growth underneath the winter snowpack. Winter-Adapted microorganisms, which had lower optimal growth temperatures and higher proteolytic potential prior to snowmelt (e.g. Figure 2b, 3g), were mainly responsible for producing N-containing monomers under the winter snowpack. The organic N monomers that accumulated prior to snowmelt were rapidly taken up and degraded for biosynthesis and energy production during the microbial bloom (Figure 4a, 5a). Synergistic coupling of organic and inorganic N transformations was observed in Winter-Adapted *Bradyrhizobia* spp. MAGs, which fermented amino acids and utilized the organic acids produced as electron donors to produce energy via denitrification during snowmelt (Figure 5c).

The fact that the DON pool became more reduced and aromatic during snowmelt suggested that turnover and recycling of microbial biomass was the primary source of N during the microbial bloom. For example, we observed marked increases in gene expression of serine and metalloendopeptidases involved in cell division, peptidoglycan synthesis and recycling (Figure 3g) that was up to 100-fold higher than gene expression for proteases that produce N monomers. Endopeptidases transform organic N through deimination-like reactions that break double bonds and produce polypeptides or polypeptide-like macromolecules. Hydrogenation reactions were also the most frequently detected transformation of DON during snowmelt (Figure 3e), consistent with electron transfer to break imine, enamine and similar double bonds in aromatic or heterocyclic DON (Figure 3g, 6c). These results imply that increased DON stocks observed during snowmelt were associated with cell division, biomass turnover and degradation of aromatic DON. Consequently, the magnitude and rate of soil N cycling overwinter should be proportional to the size of the snowmelt-associated microbial bloom.

Multiple lines of evidence suggest that distinctions between a ‘slow’ soil N cycle during winter versus a ‘fast’ soil N cycle in summer^58^ should be revisited for ecosystems that experience a seasonal snowpack. First, we observed that microorganisms that were most active underneath the snowpack also had the fastest predicted growth rates based on genome properties (Figure 2d). These results are consistent with a lab study showing that soil bacterial isolates collected during winter had faster growth rates compared to isolates enriched from summer soils^59^. Ecological theory also predicts that resource scarcity during snowmelt should lead to selection for microbial taxa with life history traits - such as fast growth rates - that maximize resource acquisition^60^. Second, we observed high gene expression for endopeptidases involved in cell division, peptidoglycan synthesis and recycling which implies high turnover of microbial biomass overwinter and during snowmelt. Third, the molecular and stoichiometric properties of the DON pool, such as becoming more reduced and aromatic, are indicators of increasing microbial processing during winter and snowmelt^61^. These results together challenge the notion of a ‘slow’ soil N pool that passes through microbial biomass only once or twice during winter^58^. In contrast, it appears that rapid microbial growth and turnover provides N to fuel the snowmelt-associated microbial bloom.

The metabolic activity of Spring-Adapted ammonia-oxidizing archaea substantially increased after snowmelt (Figure 6d) and co-occurred with a significant pulse of soil nitrate (Figure 1i). These results highlight the notable role that Spring-Adapted archaea had in increasing soil N mobilization at the start of plant growing season. Spring-Adapted ammonia-oxidizing archaea had high gene expression for urea production and consumption, but limited ability to otherwise utilize N monomers. By contrast, some Winter-Adapted organisms had high potential to mineralize N from osmolytes such as glycine betaine, or from nucleotides released from lysed microbial biomass, thereby providing additional sources of ammonia to Spring-Adapted ammonia-oxidizers. These results highlight the connected N cycling activities of Winter- and Spring-Adapted organisms after snowmelt that coupled organic and inorganic N transformations that increased N mobilization after snowmelt in spring.

We contend that nitrate produced by high nitrification activity after snowmelt was rapidly re-assimilated because we observed high gene expression for dissimilatory and assimilatory nitrate reduction to ammonia in June (Figure 6b). Recycling and re-mineralization of soil N in spring is reflective of a ‘closed’ soil N loop that provides ample N to meet plant and microbial N demand for growth during warmer periods^62^. Because the timing of high nitrate production ensued after a period of high soil-saturation that occurred during snowmelt^12,63^, it is unlikely that significant amounts of nitrate produced by Spring-Adapted organisms was transported to the subsurface by the retreating groundwater table. Rather, it appears that riverine nitrate that bears an isotopic signature derived from nitrification originates deeper in the subsurface^15,64^, or from locations adjacent to the river that experience more frequent periods of groundwater incursion or periodic river inundation (e.g. floodplain meanders) ^65–67^.

Combining multi-omic data sets had several advantages, as well as limitations, compared to previous approaches for understanding soil N cycling during and after snowmelt. For example, a previous study that used biochemical enzyme assays concluded that proteolytic activity is highest in spring after snowmelt^68^. The advantage of biochemical enzyme assays, compared to our multi-omic approach, is that biochemical enzyme assays provide a rate estimate of N monomer production. The disadvantage is that such assays represent only deamination-like reactions that produce N monomers (e.g. amino acids). In contrast, we detected thousands of DON compounds, and untargeted transformation analysis of the metabolome combined with gene expression data, allowed us to infer that serine-endopeptidase and metalloendopeptidase activity and transformations of DON that break double bonds (e.g. hydrogenation) were the major DON transformations associated with the microbial bloom. This result is important because biochemical enzyme assay data is widely used to infer rates of DON depolymerization^69–71^, but our results showed that most DON depolymerization reactions did not produce N monomers.

Our study implies that climate warming could significantly affect soil N cycling during and after snowmelt in mountain ecosystems. For example, low-to-no snow winters are expected within the next 35 to 60 years in the western U.S.^18^. Reduced winter snow cover will increase winter soil freezing and limit snowmelt infiltration as well as groundwater upwelling into surface soils during snowmelt^63^. If frozen soil limits the activity of Winter-Adapted organisms that primarily break down plant litter overwinter, then it is possible that snowmelt-associated blooms will be smaller with less soil N recycling during winter. In addition, the phenological timing of soil N cycling in spring is currently well-coupled with plants breaking winter dormancy. However, earlier snowmelt could lead to asynchrony in the timing of soil N mobilization and plant N demand. Thus, climate change may affect plant productivity and watershed N retention indirectly through climate change effects on the soil N cycling microbial community during winter and after snowmelt.

## Materials and Methods

The East River Watershed is located in Gunnison County, Colorado near the town of Crested Butte, CO (38°57.5’ N, 106°59.3’ W). Elevations within the East River Watershed range from 2750 to 4000 m. Snow cover in winter typically persists 4 to 6 months (e.g., November through May) followed by an arid summer with intermittent, monsoonal rain events that occur from July through September. Minimum annual daily air temperature occurs in January (-14 ± 3° C), whereas the maximum daily air temperature typically occurs in July (23.5 ± 3° C). Summer air temperature has increased by 0.5 ± 0.1 °C per decade since 1974^72^. Annual precipitation is approximately 1200 cm with the majority (> 80%) falling as snow during winter^2,73^. Over the past 50 years, maximum annual snow depth was 465 cm and the date of snowmelt has advanced earlier in the spring by 3.5 ± 2 days per decade^74^.

### Soil Collection and Pore Water Sampling

We sampled soils and soil pore water (i.e. soil leachate) at an upland hillslope location that was adjacent to the main stem of the East River (elevation ∼2775 m). Plant composition at this location are a mixed, montane meadow community comprised of perennial bunchgrasses (e.g., *Festuca arizonica*), forbs (e.g., *Lupinus* spp., *Potentilla gracilis, Veratrum californicum*), and shrubs (*Artemisia tridentata*)^75^. The sampling locations were encompassed by an approximately 80 m x 100 m area that was 100 m upslope from the edge of the East River riparian floodplain. The collection of soil samples has been described previously^12^ and briefly were collected on four dates starting at maximum winter snow depth (March 7, 2017), during the peak snowmelt period (May 9, 2017), during the plant growing season after the complete loss of snowpack (June 9, 2017), and lastly in autumn after plant senescence (September 16, 2017). When there was no snowpack (September and June), soil samples were collected from two locations within 6 plots and then split into three discrete depth increments; 0 to 5 cm, 5 to 15 cm, and 15 cm + below the soil surface. A ∼10 g subsample from each soil core at each depth was placed immediately on dry ice in the field, frozen and archived for metagenomics and metabolomic analysis (described below). The remainder of the soil core was allocated to physical and bio-chemical characterization (described below) and was stored at 4 °C until further analysis.

Soil temperature, soil volumetric water content, and water potential were measured continuously starting in October 2016 at 2 locations within each plot and reported previously^12^. Ceramic cup tension lysimeters were installed within each plot at 25 cm, 50 cm, and 150 cm below the surface and used to sample soil pore water in June. Low soil water content at the field site inhibited sampling the lysimeters in September (post-plant senescence) and March (winter). In May (snowmelt) we sampled soil pore water using a ¼ inch stainless steel tube inserted into the soil at discrete soil depths. Soil pore water was sampled with 60 mL gas tight syringes, filtered through 0.2 µM filters (Acrodisc, Pall Corportation), and stored in pre-evacuated 60 mL glass serum bottles fitted with 20 mm butyl-rubber septa (Bellco Glass) We sampled the tension lysimeters in June by applying a negative pressure overnight to allow soil pore water to diffuse into the lysimeter cup and sampled within 24 hours of applying the vacuum.

### Soil Chemical Properties, Microbial Biomass and N Pools

Microbial biomass N was measured using the chloroform-fumigation extraction method^76^. A 5 g subsample of each soil (field-moist) was extracted in 25 mL of 0.5M K_2_SO_4_ on an orbital shaker table for 60 min, then gravity filtered through pre-leached #42 Whatman filter paper and frozen until further analysis. A separate 5 g subsample was fumigated with ethanol-free chloroform for 7 days and then extracted as stated above. Microbial biomass N was quantified as total dissolved N measured in the fumigated minus non-fumigated extracts^72^. Total dissolved N was measured colorimetrically as nitrate (NO_3_-N) after alkaline persulfate oxidation using a Versamax microplate spectrophotometer (Molecular Devices). We did not apply an extraction correction factors for incomplete microbial biomass lysis during the fumigation.

Extractable inorganic N, total free primary amines, and DON pool sizes were also measured in the unfumigated soil extracts. Dissolved inorganic N was measured colorimetrically using a Versamax microplate reader (Molecular Devices) as ammonium (NH_4_-N) + nitrate (NO_3_-N) in both soil extracts and soil pore water samples^77,78^. We estimated DON as the difference in total dissolved N minus inorganic N. Lastly, total free primary amine N was measured fluorometrically using the o-phthaldialdehyde — β-mercaptoethanol (OPAME) method^79^ and a SpectraMax 340PC fluorescent microplate reader (Molecular Devices). All concentrations measured in the soil extracts were corrected for the field-moist water content of the soil.

### Nucleic Acid Co-Extractions

DNA and RNA were co-extracted from 5 to 7 technical replicates of each soil sample on ice by adding 0.5 mL phenol:chloroform:isoamyl alcohol (25:24:1) (Sigma-Aldrich, St. Louis, MO, USA) to 0.5 g of soil in a 2 ml Lysing Matrix E tube (MP Biomedicals, Solon, OH, USA), followed by addition of 0.5 mL of CTAB buffer (5% CTAB, 0.25M phosphate buffer pH 8.0, 0.3M NaCl) and 50 μL of 0.1M aluminum ammonium sulfate. The samples were homogenized at 5.5 m/s for 30 s in a FastPrep-24 instrument (MP Biomedicals, Solon, OH, USA), then centrifuged at 16K g for 5 min at 4 °C. The aqueous phase was removed and transferred to MaxTract High Density 2 mL tubes (Qiagen Inc, Valencia, CA, USA). Samples were then extracted a second time as described above and the aqueous phase from technical replicates for each soil sample were combined. Sodium acetate (3M sodium acetate, 1/10^th^ volume of total aqueous phase) and ice-cold ethanol (100%, 2X volume of total aqueous phase) were added, the samples were vortexed briefly, and a crude nucleic acid pellet was precipitated overnight at -20 °C.

Separation of DNA and RNA was completed using the AllPrep DNA/RNA Mini Kit (Qiagen Inc, Valencia, CA, USA). The amount of DNA or RNA extracted was quantified using the Qubit 1X dsDNA Broad Range Kit or Qubit RNA High Sensitivity Kit, respectively (ThermoFisher Scientific). DNA and RNA quality were assessed using a 2100 Bioanalyzer instrument (Agilent Technologies, Santa Clara, USA). DNA and RNA were stored separately at - 80 °C prior to metagenome and metatranscriptome analyses (described below).

### FT-ICR MS and ^1^H NMR Characterization of Dissolved Organic Matter

Metabolomic data were collected at the Department of Energy’s Environmental Molecular Sciences Laboratory (EMSL; Richland, WA). Extractable dissolved organic matter (DOM) was extracted from 2 g subsamples of freeze-dried soil as described previously^77,78^. A high-speed ball mill was used to homogenize the subsamples prior to extraction. We first added 1.5 mL MilliQ water (H_2_O) to the soil, extracted the H_2_O-soluble fraction by shaking for 2 hours at room temperature, centrifuged the soil slurry at 15,000 x g for 30 min to pellet the soil, and lastly decanted the supernatant to collect the H_2_O-soluble fraction. Soil pore water samples collected for metabolomics analysis were filtered through 0.2 µM filters (Acrodisc, Pall Corporation) in the field and subsequently frozen prior to preparation for FT-ICR MS or ^1^H-NMR analysis (described below).

Molecular properties of DOM were characterized by electrospray ionization (ESI) using a 21T FT-ICR MS^80^. LC-MS grade methanol was used to dilute the H_2_O-soluble (1:2 v/v) fraction prior to injection to assist in ionization. Three technical replicates were run for each sample, samples were randomized, and an autosampler was used for direct infusion. Experimental conditions were optimized for this study. Briefly, a standard Thermo Heated-ESI (HESI) source was used to generate negatively charged molecular ions. Samples were introduced to the HESI source with a fused silica tube (50 µm i.d.) using an Agilent 1200 series pump (Agilent Technologies) at a flow rate of 4 µL min^-1^. Experimental conditions were as follows: scan average was set at 100µS; needle voltage +3.4 kV; S-Lens RF level set to 60%; source heater temperature operated at 40 °C and the heated resistively coated glass capillary operated at 275 °C.

Data was collected over 22 minutes with seven of the 100µS scans averaged for each sample and calibrated using an organic matter homologous series separated by 14 Da (-CH_2_ groups) and used to convert raw spectra to a list of *m/z* values by applying a signal to noise (S/N) ratio > 2, mass measurement error < 1 ppm, an absolute intensity threshold value of 100 taking into account the presence of C, H, N, O, P, S while excluding other elements. Features (i.e. *m/z* values) were kept if detected in at least two of three technical replicates. Chemical formulae were assigned using the Compound Identification Algorithm described previously and implemented by the software Formularity^81^.

We used the online software FT-MS R Exploratory Data Analysis (FREDA; https://shinyproxy.emsl.pnnl.gov/app/freda) to calculate the C to N ratio, estimate double bond equivalents (DBE), the nominal oxidation state of carbon (NOSC), and the aromaticity for each compound found in each sample. All compounds were categorized into biochemical compound categories (e.g. ‘phyto-oxy-aromatic’, ‘peptide’, ‘amino sugar’, ‘nucleotides’ or ‘unmatched’ based on multiple elemental stoichiometric classification rules as described previously^82^. Transformations of the DON pool was inferred by calculating the pairwise mass difference between all compounds detected in each sample and comparing those mass differences to a reference list of 1255 precise masses of commonly observed biochemical transformations^83,84^. The relative abundance for each transformation in each sample was calculated by dividing the sum of each transformation by the total number of transformation detected in each sample. We also grouped transformations into four categories (N-containing, P-containing, S-containing, or CHO-only, or OH-only) following methods described previously^85^. For example, transformations that involved chemical moieties containing N (e.g. amino acids) were categorized as ‘N-containing’, whereas transformations with chemical moieties comprised of carbon (C), oxygen (O) and hydrogen (H) were categorized as ‘CHO-only’. Transformations that contained both N and either phosphorus (P) or sulfur (S) were grouped with the N-containing category.

^1^H-NMR metabolite characterization of field-present metabolites was performed on the same extract as used for FT-ICR MS (described above). The soil water extracts were diluted 10% (v/v) with 5 mM 2,2-dimethyl-2-silapentane-5-sulfonate-d6 (DSS-d6) as an internal standard, thoroughly mixed then transferred to 3-mm NMR tubes. All NMR spectra were acquired on a Varian 600 MHz VNMRS spectrometer equipped with a 5 mm triple resonance salt-tolerant cold probe and a cold-carbon pre-amplifier at a regulated temperature of 298K. The 90° ^1^H pulse was calibrated prior to the measurement of each sample. The one-dimensional (1D) ^1^H spectra were acquired using a nuclear Overhauser effect spectroscopy (NOESY) pulse sequence with a spectral width of 12 ppm and 512 transients. The NOESY mixing time was 100 ms, and the acquisition time was 4 s, followed by a relaxation delay of 1.5 s during which presaturation of the water signal was applied. Time-domain free induction decays (57,472 total points) were zero filled to 131,072 total points prior to Fourier transform. Chemical shifts were referenced to the ^1^H methyl signal in DSS-d6 at 0 ppm. The 1D ^1^H spectra were manually processed, assigned metabolite identification, and quantified using Chenomx NMR Suite 8.3. Metabolite identification was based on matching the chemical shift, J-coupling, and the intensity of experimental signals to compound signals in the Chenomx, HMDB and custom in-house databases. Quantification was based on fitted metabolite signals relative to the internal standard (DSS-d6). Concentrations were normalized by soil mass and H_2_O volume used for extraction to units of umol / g. Signal-to-noise ratios (S/N) were measured using MestReNova 14 with the limit of quantification equal to an S/N of 10 and the limit of detection equal to an S/N of 3. Standard 2D experiments such as ^1^H /^13^C - heteronuclear correlation (HSQC) or 2D ^1^H /^1^H Total Correlation spectroscopy (TOCSY) further aided corroboration of several metabolite identifications where there was sufficient S/N.

### Metagenome and Metatranscriptome Library Preparation and Sequencing

Metagenome and metatranscriptome sequencing libraries were generated (mean insert size 255 bp and 215 bp for DNA and RNA, respectively) and sequenced at the U.S. Department of Energy’s Joint Genome Institute (JGI, Berkeley, CA USA) using the Kapa Library with Real Time PCR compatible with Illumina series (Kapa Biosystems, United Kingdom). For metagenomes, high molecular weight DNA was sheared using a Covaris LE220 focused-ultra sonicator and size selected using double SPRI. Then size selected fragments were end-repaired and ligated with adapters containing molecular index barcodes unique to each sequencing library. For metatranscriptome libraries, rRNA was first depleted using the Illumina’s Ribo-Zero rRNA Removal Kit (Bacteria), and stranded cDNA libraries were generated using the Illumina TruSeq Stranded Total RNA kit. The rRNA-depleted RNA was fragmented and reversed transcribed using random hexamers and SSII followed by second strand synthesis. The fragmented cDNA was treated with end-pair, A-tailing, adapter ligation, and 10 cycles of PCR. Metagenome and metatranscriptome libraries were each quantified separately using a Roche LightCycler 480 real-time PCR instrument. Sequencing of metagenome and metatranscriptome libraries were completed using the Illumina NovaSeq sequencer with the NovaSeq XP V1 reagent kits, S4 flowcell, following a 2×151 indexed run recipe. Paired metagenome and metatranscriptome data are available under GOLD Study ID Gs0135149 (https://gold.jgi.doe.gov/study?id=Gs0135149). Sequencing yielded 25.6 ± 5.8 Gb (range: 18.3 – 46.2 Gb) of DNA sequence and 26.2 ± 5.68 Gb (range: 5.1 - 37.6 Gb) of RNA sequence per sample.

### Metagenome Assembly, Binning and Annotation

Raw metagenome reads were initially assessed for quality using procedures described previously^86^ with processing scripts available and described in “Filtered_Raw_Data” folders for GOLD Study ID Gs0135149 (https://gold.jgi.doe.gov/study?id=Gs0135149). Reads associated with potential contaminants (human, cat, dog, mouse references and common microbial contaminants) were removed, adapter-trimmed and further trimmed from where quality dropped to 0 using BBDuk (version 38.08 and 38.24 for DNA and RNA respectively). Post-trimming, BBDuk was also used to remove reads that satisfied any of the following filters: (1) contains 4 or more undetermined bases (‘N’), (2) average quality score across the read smaller than 3, (3) minimum length <= 51 bp or 33% of the full read length. For RNA reads, the additional filtering included removal of reads from ribosomal RNA (rRNA) through detection of common ribosomal kmers in the reads using BBDuk. Quality control filtering resulted in removal of an average of 1.3% of reads per or 1.99% of bases per sample for DNA reads. For RNA, including removal of reads of rRNA origin, quality control filtering resulted in removal of an average of 67.5 ± 24.3% (range: 3.2 – 93.3 %) of reads per sample representing an average of 68 ± 23.8% (range: 6.3 – 93.4%) of bases per sample, largely the result of rRNA read removal (99.14% per sample of the removed reads on average).

Trimmed and filtered paired-end reads were read corrected using bfc (version r181, https://github.com/lh3/bfc)^87^ with “bfc -1 -s 10g -k 21 -t 10”. Singletons with no mate pairs post filtering were removed. Corrected reads were assembled with metaSPAdes assembler for each sample (version 3.11.1)^88^ using the following range of k-mers as follows: “spades.py -m 2000 -- only-assembler -k 33,55,77,99,127 --meta -t 32”. To determine read representation in each assembly, the filtered read set from the sample was mapped to the assembly and coverage was inferred using bbmap (version 37.78) with default parameters except ambiguous=random. Overall across 48 samples, 36.38 ± 8.3 % (range: 19.9 – 55.2%) of the filtered DNA reads mapped back to the sample specific assemblies. Full statistics describing quality control filtering and assembly statistics is given in Supplemental Table 9.

Metagenome binning was performed in ggKbase (https://ggkbase.berkeley.edu/) for each assembly. Contigs longer than 1 kb were binned using the automated binning algorithms MetaBat^89^, Maxbin2^90^, and Concoct (version 1.1.0)^91^. DASTool was used to select the best set of bins from each metagenome sample based on dereplication, aggregation and scoring^92^. This resulted in 522 metagenome-assembled genomes (MAGs). MAGs were checked for completeness against a set of 51 bacterial single copy genes or a set of 38 archaeal single-copy genes. A subset of 167 MAGs were categorized as ‘high-quality’ which passed quality filtering in checkM^93^ with more than 75% completeness and less than 25% contamination. Contigs and predicted orfs were annotated using the IMG annotation pipeline (v4.16.4)^94^ and MAG trait inference were performed using the microTrait pipeline^95^.

For mapping RNA reads to MAGs, quality control filtered RNA reads were mapped separately in two ways: (1) RNA reads were mapped to checkM filtered MAGs from all samples, and (2) RNA reads were mapped to all orfs predicted from all assembled contigs regardless of read binning status. Mapping was performed with bbmap where multiple mapping locations were resolved randomly (ambiguous=random). RPKM values as calculated by bbmap were used for further analysis.

### Statistics

All statistical analyses were completed using R v4.0.0^96^. We tested for differences in extractable soil N pools across time, soil depths and their interaction using a two-factor analysis of variance using the R package ‘rstatix’^97^, verifying assumptions of normality and equal variance by inspecting quantile and residual plots ‘ggResidual’^98^. The degree to which metagenome-assembled genome (MAG) abundance or activity varied across sampling dates or soil depths was assessed by permutational multivariate analysis of variance (perMANOVA, permutations n = 999) based on Bray-Curtis dissimilarity distances calculated using the R package ‘vegan’^99^. MAG abundance was inferred from genome coverage and activity was inferred from total gene expression mapped to each MAG. Pairwise comparisons of community dissimilarity across months or depths were determined using the R package ‘mctools’^100^.

Hierarchical clustering was used to classify MAGs into ecological response groups according to whether maximum MAG activity or abundance occurred under the snowpack in winter versus during the snowmelt-period or after snowpack loss in June. The average-linkage method was used to calculate distances between clusters and dendrograms were visualized using ‘ggdendro’^101^. We used the non-parametric Kruskal-Wallis test from the ‘rstatix’ package to determine whether microbial N cycling traits or N-containing small molecule metabolite concentrations were different across sampling dates. In the case of N cycling trait expression, p-values were adjusted for multiple comparisons by the Benjamini and Hochberg false discovery rate (‘fdr’) method using the R package ‘stats’. Figures were made using the ‘ggplot2’ and ‘patchwork’ R packages^102,103^, as well as using Biorender.com

## Data Availability

The raw shotgun metagenome data has been deposited and is available through NCBI’s SRA and BioSample repository under umbrella project PRJNA116881. Associated metadata is available through JGI’s Genomes OnLine (GOLD) system under GOLD Study ID Gs0135149 (https://gold.jgi.doe.gov/study?id=Gs0135149). Sample metadata, individual metagenome assemblies, and metatranscriptome data are available through the National Microbiome Data Collaborative, along with links to the NCBI BioSample identifiers at: https://data.microbiomedata.org/details/study/nmdc:sty-11-dcqce727 and ESS-Dive (doi:10.21952/WTR/1573029).

## Acknowledgements

We thank Mark Conrad, Rosemary Carroll, and Wendy Brown for their critical efforts in digging snow pits that enabled soil sampling during winter. We also wish to thank Jenny Reithel and Shannon Sprott at Rocky Mountain Biological Laboratory (RMBL) for their general assistance at the field site. Rocky Mountain Biological Laboratory (RMBL) lab spaces and equipment are funded in part by the RMBL Equipment (Understanding Genetic Mechanisms) Grant, DBI-1315705. (A portion of) This research was performed under the Facilities Integrating Collaborations for User Science (FICUS) program (proposal: 10.46936/fics.proj.2019.50964/60000127) and used resources at the DOE Joint Genome Institute (https://ror.org/04xm1d337) and the Environmental Molecular Sciences Laboratory (https://ror.org/04rc0xn13), which are DOE Office of Science User Facilities operated under Contract Nos. DE-AC02-05CH11231 (JGI) and DE-AC05-76RL01830 (EMSL). This material is based upon work supported as part of the Watershed Function Scientific Focus Area at Lawrence Berkeley National Laboratory funded by the U.S. Department of Energy, Office of Science, Office of Biological and Environmental Research under Award Number DE-AC02-05CH11231.

