## Supplemental Figures for "Coordinated organic and inorganic nitrogen transformations fuel soil microbial blooms and increase nitrogen retention during snowmelt"

Supplemental Figure Captions

**Supplemental Figure 1** Non metric dimensional scaling of (a) variation in genome cover or (b) variation in genome expression across sampling dates using bray-curtis distance. Permutational analysis of variance was used to test for the effects of month of sampling, depth and their interaction and are shown in the figure.

**Supplemental Figure 2** Hierarchical clustering was used to categorize MAGs based on the date each MAG was most abundant based on genome coverage across time.

**Supplemental Figure 3** Heatmap showing 103 transformations of organic N that produced monomers

**Supplemental Figure 4** Gene expression for genes involved in fermentation of branched-chain amino acids. Gene expression was normalized by z-scoring gene expression across time.

**Supplemental Figure 5** Gene expression for genes involved betaine degradation. Gene expression was normalized by z-scoring gene expression across time.

**Supplemental Figure 1**

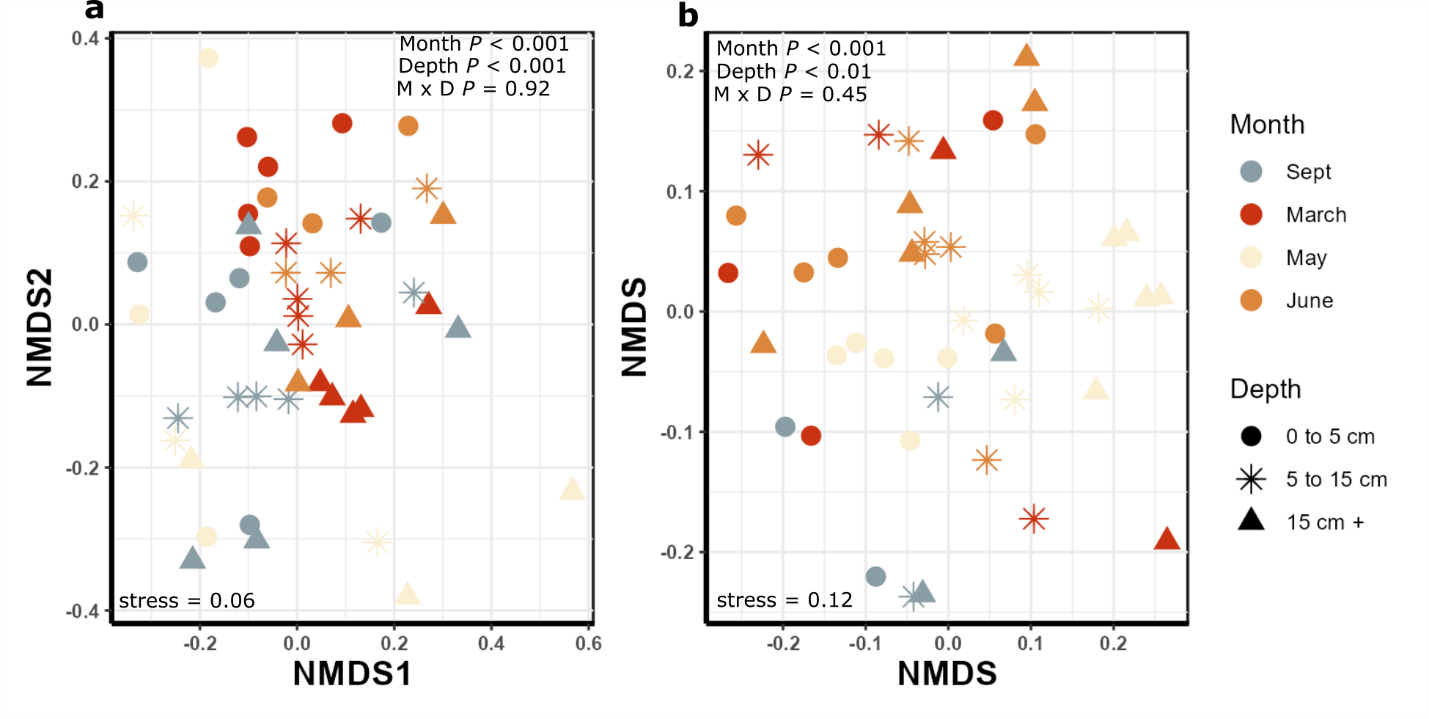

**Supplemental Figure 2**

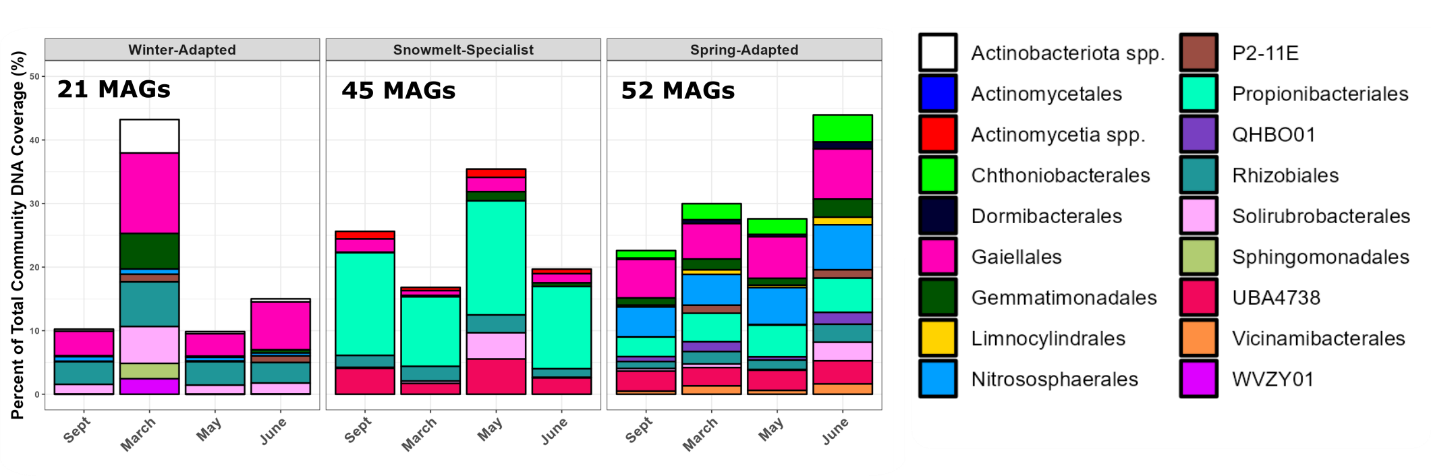

**Supplemental Figure 4**

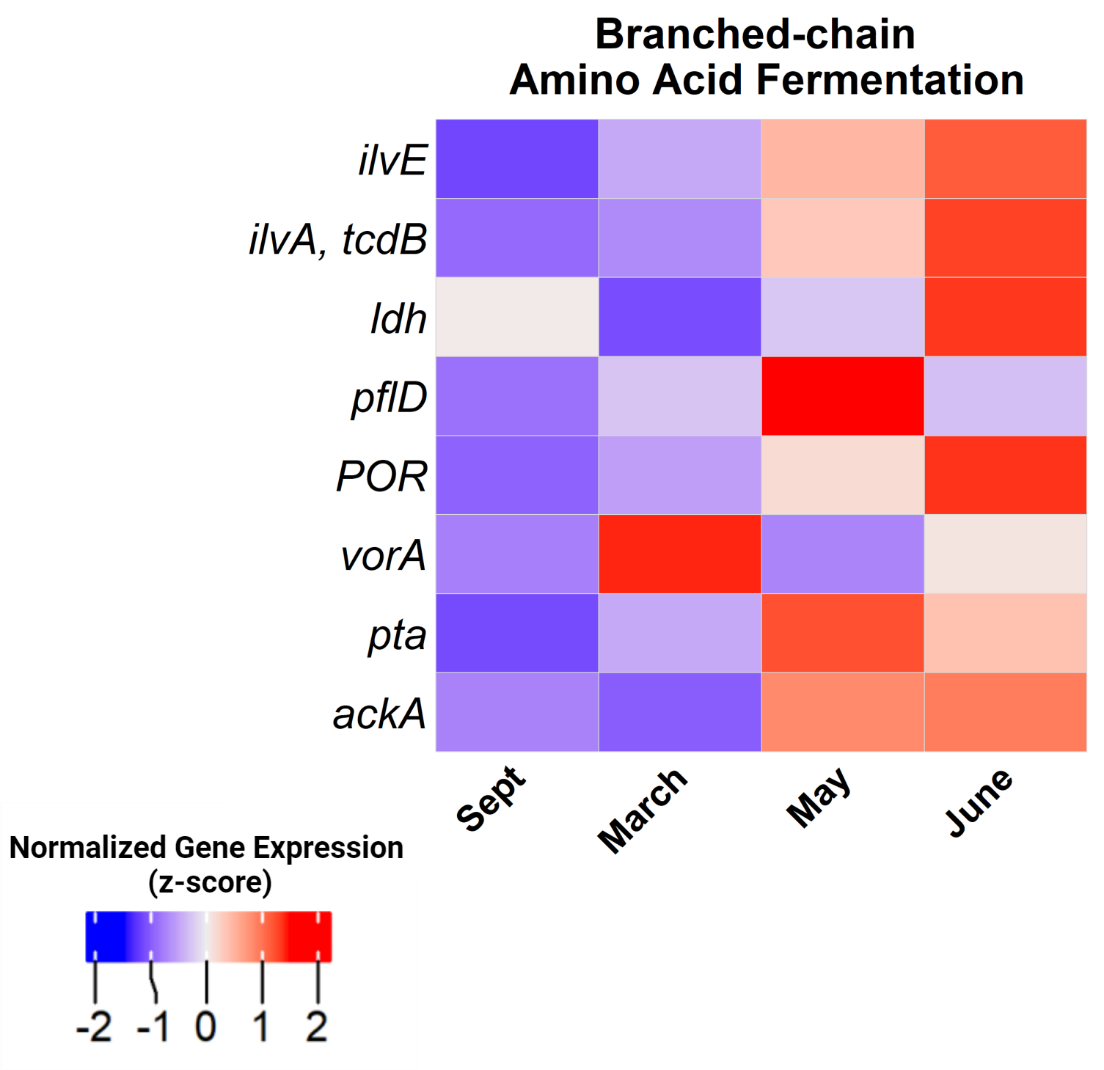

**Supplemental Figure 5**

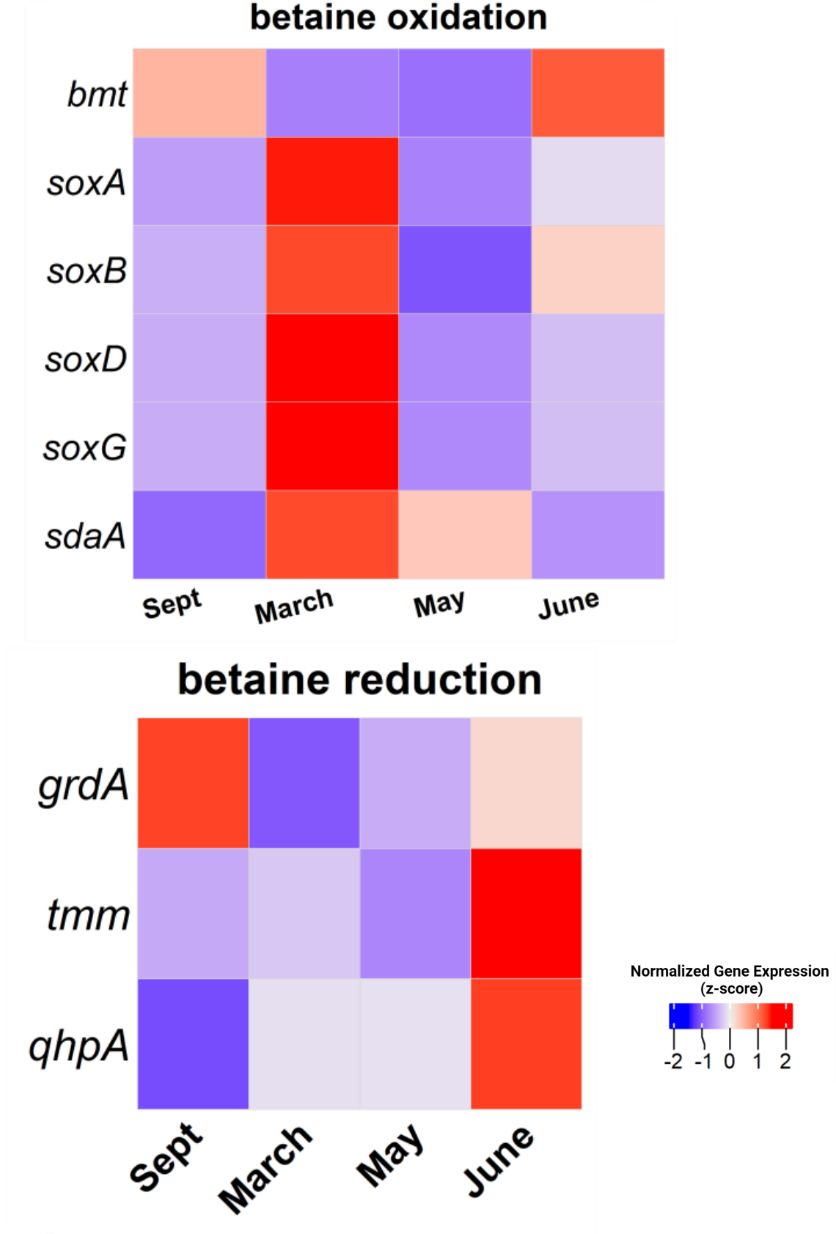
